## Supplemental Figure 1 for "A novel HIV triple broadly neutralizing antibody (bNAb) combination-based passive immunization of infant rhesus macaques achieves durable protective plasma neutralization levels and mediates anti-viral effector functions"

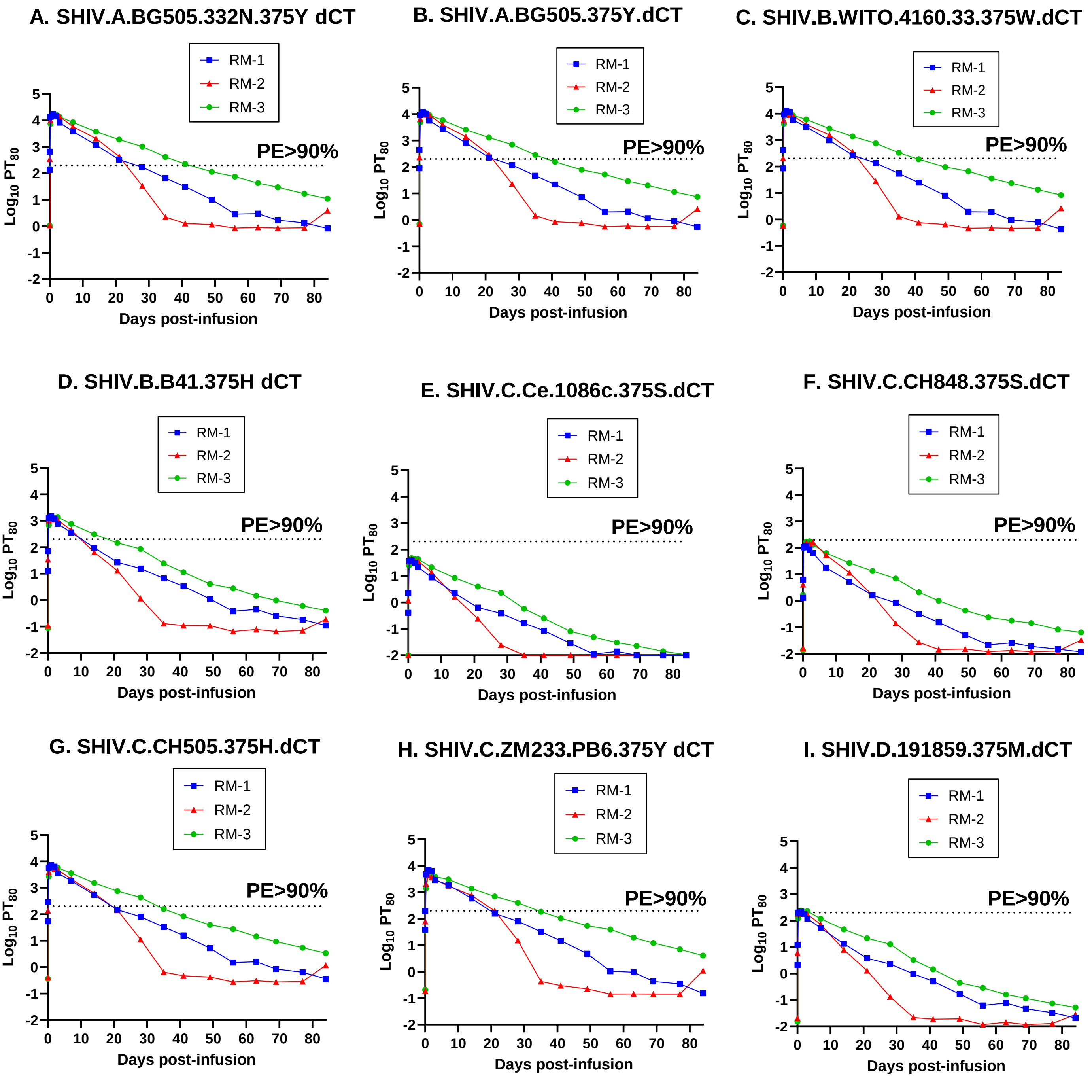


**Figure S1. Predicted preventative efficacy (PE) of the triple bNAb combination against cross-clade SHIV variants.** $PT_{80}$ values of Rh bNAb combination against **(A)** SHIV.A.BG505.332N.375Y.dCT **(B)** SHIV.A.BG505.375Y.dCT **(C)** SHIV.B.WITO.4160.33.375W.dCT **(D)** SHIV.B.B41.375H.dCT **(E)** SHIV.C.Ce.1086c.375S.dCT **(F)** SHIV.C.CH848.375S.dCT **(G)** SHIV.C.CH505.375H.dCT **(H)** SHIV.C.ZM233.PB6.375Y.dCT and **(I)** SHIV.D.191859.375M.dCT. Dashed line represents $PT_{80}$>200, equivalent to ≥90% PE of the antibody combination against the SHIV variants in rhesus macaques.
